# *S*timulus-modulated *a*pproach to *s*teady-*s*tate: A new paradigm for event-related fMRI

**DOI:** 10.1101/2024.09.20.613944

**Authors:** Renil Mathew, Amr Eed, L. Martyn Klassen, Omer Oran, Stefan Everling, Ravi S Menon

## Abstract

Functional MRI (fMRI) studies discard the initial volumes acquired during the approach of the magnetization to its steady-state value. Here, we leverage the higher temporal signal-to-noise ratio (tSNR) of these initial volumes to increase the sensitivity of event-related fMRI experiments. To do this, we introduce Acquisition Free Periods (AFPs) that allow for the full recovery of the magnetization, followed by task or baseline acquisition blocks (AB) of fMRI volumes. An appropriately placed stimulus in the AFP produces a Blood Oxygenation-Level-Dependent (BOLD) response that peaks during the initial high tSNR phase of the AB, yielding up to a ∼50% reduction in the number of trials needed to achieve a given statistical threshold relative to conventional fMRI. The silent AFP can be exploited for the presentation of auditory stimuli or uncontaminated electrophysiological recording and its variable duration allows aperiodic stimulus or response-locked signal averaging as well as gating to physiology or motion.

---

BOLD fMRI has been a mainstay of non-invasive functional brain imaging for over 30 years^1–3^. fMRI experiments rely almost exclusively on versions of the echo-planar imaging (EPI) sequence to map resting-state or stimulus-evoked BOLD signal changes over time^4,5^. Typically, a dozen or so initial volumes, acquired during the approach to steady-state equilibrium of the magnetization are either discarded or treated as dummy scans. Unfortunately, these are the images with the highest SNR. The approach to steady-state of the magnetization is described by the Bloch equations^6^, is typically 5 – 10 s, and is influenced by the radio frequency pulse flip angle (FA) and the longitudinal relaxation time constant (T1) of the tissue. Since T1 increases with field strength, the approach presented in this manuscript has particular benefit at ultra-high magnetic fields. While generally considered as an annoyance for paradigm design, stimulus triggering or analysis pipelines, EPI volumes acquired during the approach to equilibrium can offer valuable insights into tissue-specific relaxation parameters and have been proposed for improving grey/white matter tissue discrimination, resulting in a reduction of false positives in event-related activation maps^7^.

In this study, we demonstrate an event-related fMRI method that allows stimuli to be presented and/or physiological signals to be monitored during a quiet period (AFP) with no radio frequency or gradients from the MRI process. This has substantial advantages for auditory processing experiments, cognitive tasks requiring high levels of concentration or triggering to asynchronous phenomena such as eye movements, phase of a gamma rhythm or respiration. Furthermore, this MRI acquisition-free period allows high-fidelity recording of stimulus responses using EEG or single-unit recording before reading out the stimulus-evoked BOLD signal. The AFP length can vary from trial-to-trial but must have a minimum duration of ∼5-10 s depending on the grey matter T1. The duration of the AFP might suggest that this approach is not as time-efficient as the conventional steady-state approach where images are acquired continuously, however we capitalize on the substantially higher magnetization available during the initial decay period to achieve equal or enhanced stimulus-evoked BOLD signal detection relative to a conventional steady-state approach. In our implementation, recurring AFPs with no gradients or radio frequency pulses present allow for the full recovery of the magnetization and are followed by ABs consisting of multiple rapidly-acquired volumes of a typical fMRI sequence such as EPI. The EPI volumes acquired have an initially higher SNR that decays to the usual steady-state value. Incorporating this into an fMRI paradigm, we structure ABs into identical length independent baseline and task ABs. To achieve optimum signal detection sensitivity for the stimulus, we present the stimulus such that the peak of the hemodynamic response that underlies the BOLD signal is captured in task AB volumes during the higher SNR phase of the magnetization decay. Because the *s*timulus is modulating the *a*pproach to *s*teady-*s*tate, we abbreviate this paradigm as SASS. Here we demonstrate the advantages of using the SASS paradigm and compare the detection sensitivity with the traditionally used steady-state (SS) EPI sequence, keeping either the experiment time or number of trials constant. Experiments were conducted on awake common marmosets (*Callithrix jacchus*) at 9.4 T using visual and auditory stimuli and on humans at 3 T using visual stimuli. Utilizing these broadly available high- and ultra-high-field systems, we examined how the initial magnetization level and decay in an AB influence tSNR improvements across field strengths and different noise limiting regimes.

## RESULTS

### Acquisition block and tSNR calculations

The basic building block for SASS paradigm consists of two elements, an AFP followed by an AB. This building block is repeated multiple times with (task AB) or without (baseline AB) a presented stimulus (Fig.1a) (Methods). Within an AB, apparent transverse relaxation time (T2*)-weighted EPI (i.e. BOLD sensitive) volumes with a given repetition time (TR) and flip angle (FA) result in a magnetization that decays to a steady-state value governed by the T1 relaxation in each voxel. To characterize the noise in each voxel over the course of the AB, we computed individual voxel-wise tSNR maps for each volume within an AB (Fig.1b) because tSNR (rather than raw SNR) is an important determinant of the ability to detect BOLD signal over the noise. For our initial demonstration at 9.4 T, the grey matter tSNR maps across the AB (block length = 16 s, TR = 1 s) demonstrated a higher initial tSNR before reaching the steady-state tSNR value after 8-10 seconds (∼4-5 times the T1 constant of grey matter^8^) (Fig.1c). As a simple tissue discrimination approach, the tSNR maps were segmented using Independent Component Analysis (ICA) into grey and white matter voxels from the raw timeseries of acquired functional volumes, where the inherent difference in their T1 relaxation decays can be used for tissue discrimination (Extended Data Fig.3a, b). We observed that the grey matter tSNR of the first volume was 61% higher than the last volume in the AB (where the steady-state has been achieved) (Fig.1d). Averaged over the first four volumes (4 s) where we would place the peak of the BOLD response, the tSNR gain was in the range of 22% to 36% at 9.4 T depending on HRF characteristics, and will be dependent on T1 and TR. The higher tSNR in the initial AB volumes gave us insight into how a stimulus response captured during these initial volumes can be detected with higher signal sensitivity compared to a steady-state condition.

**Fig. 1.**
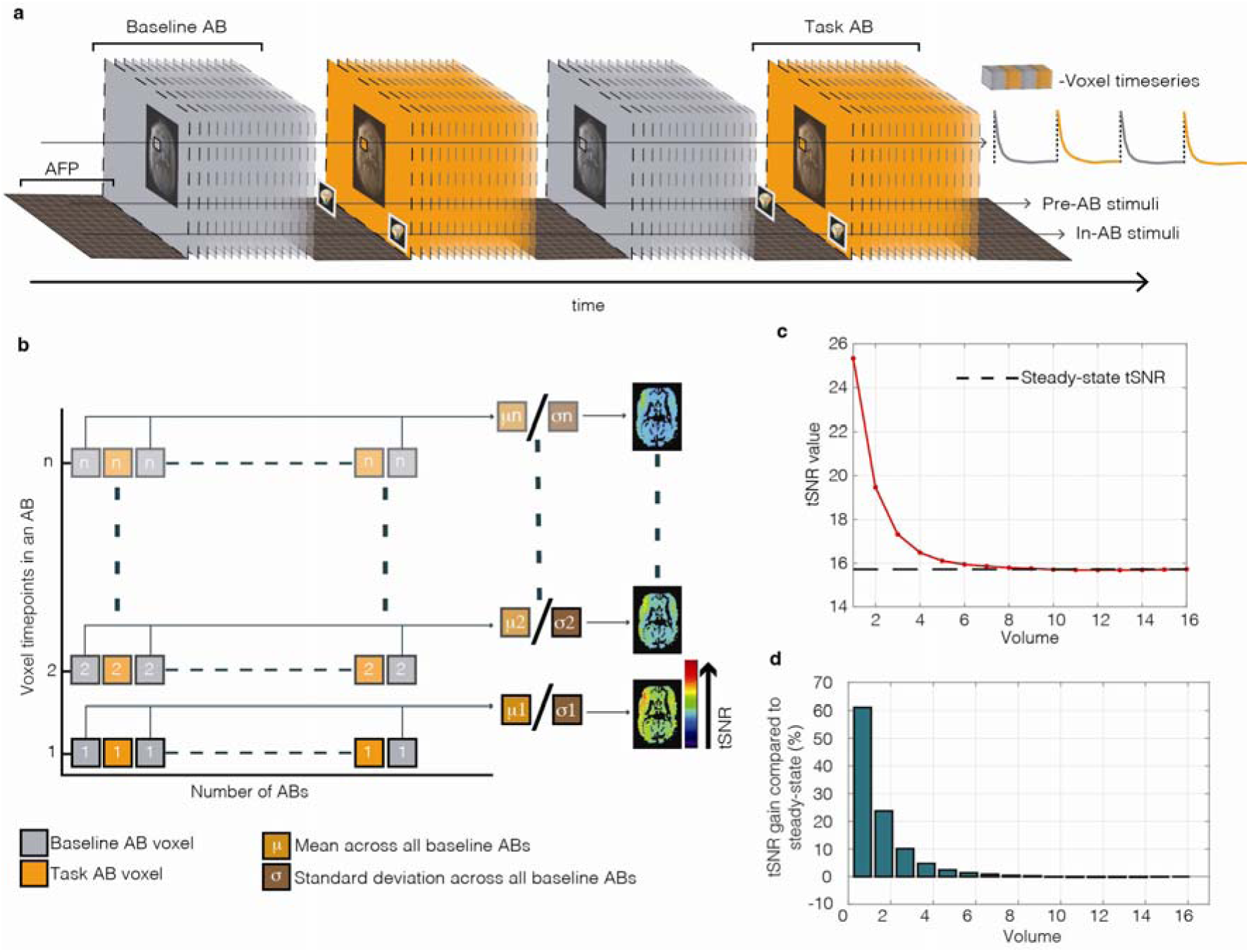
Acquisition blocks schematics and tSNR calculations at 9.4 T. **a,** Schematics of basic recurring unit in an SASS paradigm with ABs characterized into baseline and task blocks separated by AFPs. Schematics also represent pre-AB and in-AB stimulus timings used in the study. **b,** Schematics for calculating baseline mean and standard deviation from baseline ABs, used for baseline z-score normalization and tSNR calculation (tSNR calculated from all ABs). **c,** Mean grey matter tSNR variance across AB volumes with maximum value in the first volume before approaching steady-State value in the last few volumes. **d,** Bar graph showing the tSNR gam in percentage relative to last volume in steady-state.

The SASS paradigm was used to detect a stimulus-evoked BOLD response by separating the ABs into alternating identical-length task and baseline blocks. The stimulus was presented such that the subsequent BOLD response was captured during a task AB (Fig.1a). Fig.1a shows two different stimulus timings used in the study, with a stimulus presented either during the AFP immediately before the task AB (pre-AB stimulus) or within the first volume of the task AB (in-AB stimulus). To distinguish the stimulus-evoked BOLD response from the decaying magnetization signal while maintaining the fidelity of the BOLD response, we used the baseline ABs from each preprocessed fMRI run to calculate the mean and standard deviation of each voxel timepoint across all baseline ABs (Fig.1b). The baseline values were used to z-score normalize all volumes in all ABs, thereby representing each voxel timepoint within a task or baseline AB in terms of the standard deviation from their respective baseline AB mean values (Equation 2 - Methods). This volume-by-volume z-score normalization process is necessary because the imaging signal varies in time, unlike the conventional steady-state case where it is constant. The resultant z-score transformed runs are free from the influence of T1 decay, maintain the shape fidelity of the BOLD response and can be used for further statistical analysis with conventional fMRI methods. Fig.2a shows the mean raw timeseries of active voxels from a single fMRI run, reflecting the characteristic T1 relaxation of each AB within the run and Fig.2b shows the baseline z-score normalized timeseries of the same.

**Fig. 2.**
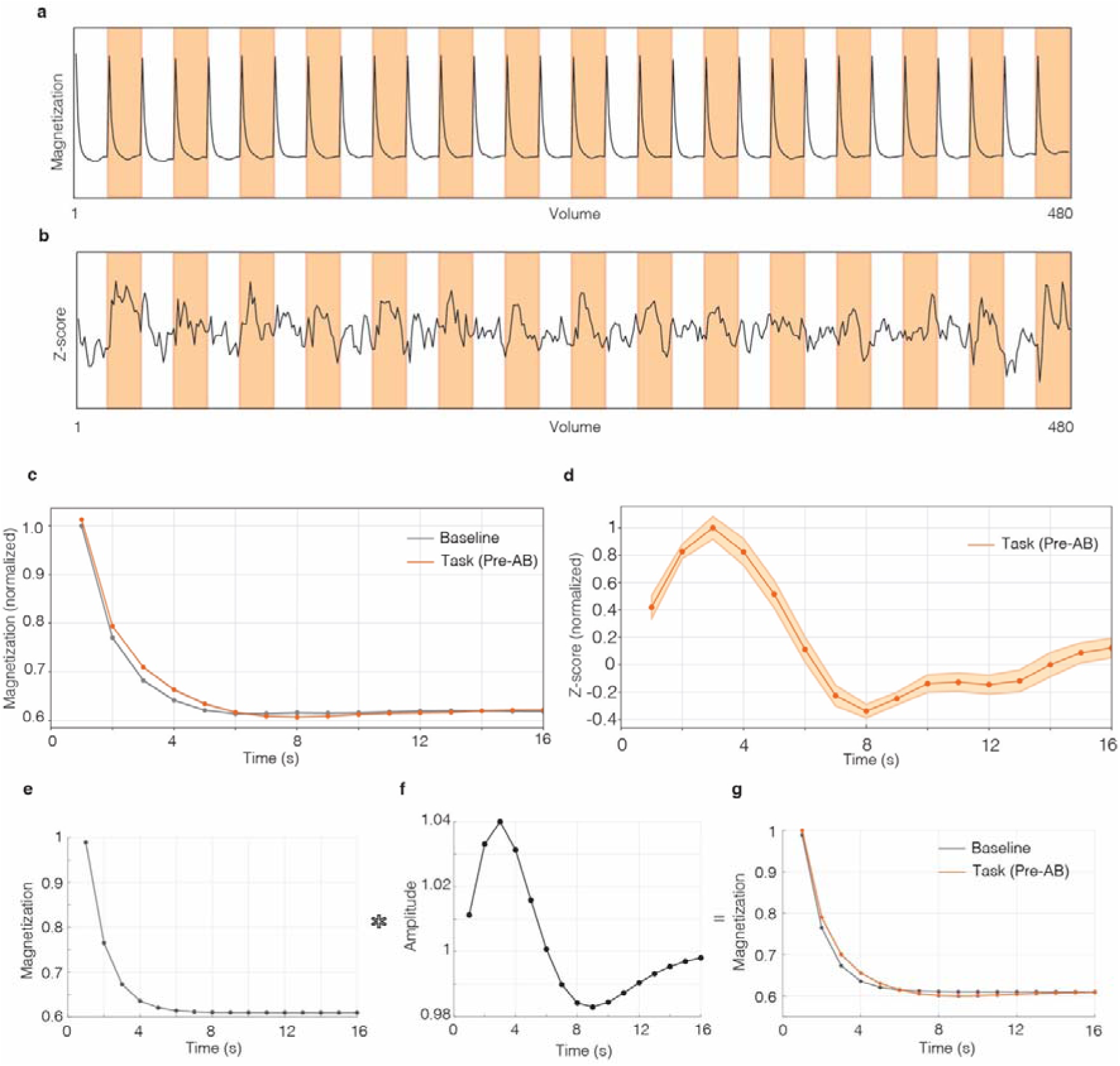
Z-score normalization and comparison of raw data to theoretical modeling for 9.4 T. **a,** Raw mean timeseries of task active voxels from a run showing characteristic magnetization decays in all ABs. task AB’s in orange color (number of volumes =480, TR 1 s). **b,** Mean z-score timeseries of task active voxels from one run after baseline z-score normalization. **c,** Mean magnetization decays from baseline and task ABs of task active voxels (for pre-AB stimuli) from a single fMRI run, both decays normalized to the maximum value of mean task AB with shaded error bars. **d,** Mean of z-score normalized task ABs of task active voxels from a fMRI run. normalized to maximum with shaded error bars. **e,** Simulated baseline magnetization decay from Bloch’s equations for TR:1 s, T1:2 2 s and FA 50 degrees, normalized to maximum of task magnetization. **f,** Truncated stimulus-HRF peaking at 3 second (marmoset monkeys) for a 1 second pre-AB stimulus (maximum BOLD percent signal change : 4%). **g,** Simulated baseline magnetization plotted with task magnetization (baseline magnetization X stimulus-HRF).

### Comparison of experimental data to theoretical modeling for 9.4 T

We compared the acquired raw data using the SASS paradigm with theoretical predictions using a series solution of the Bloch equations for the magnetization behaviour in an AB (Equation 1 – Methods). The mean task and baseline magnetization decay of active voxels for a single run containing 15 baseline ABs and 15 task ABs was computed (Fig.2c). The task was a 1 s presentation of a visual stimulus in the AFP immediately before the start of each task AB acquisition (pre-AB stimulus). A clear difference between the baseline and stimulus-evoked magnetization decay curves is observed. In a conventional steady-state fMRI experiment, the time-varying percent signal change curve accurately mirrors the shape of the BOLD response because the baseline is constant. However, in the case where the magnetization is decaying in time, the shape of the time-varying percent signal change is distorted, and the SNR also varies in time. To correct for these factors, we z-score normalized all AB voxels using the baseline ABs to estimate the temporal variance as described above. This baseline z-score normalized signal correctly reflects the shape of the BOLD response for a pre-AB stimulus, allowing for standard fMRI processing tools to be used in further analysis (Fig. 2d).

We computed the tissue magnetization decay within a baseline AB using the solution to the Bloch equations (Equation 1 - Methods) (Fig.2e). We modeled the decay with T1 = 2.2 s, FA = 50 degrees (to match the nominal FA of the experiment determined by the Ernst angle) and TR = 1 s. We then further modeled the T2*-weighted BOLD response to a 1 s pre-AB visual stimulus by convolving a standard double-gamma hemodynamic response function (HRF) peaking at 3 s with a 1 s stimulus (Fig.2f). By multiplying the calculated baseline magnetization with the resulting BOLD response^9^ (truncated stimulus-HRF to match pre-AB stimulus onset) for every volume in the AB, we generated an estimate of the stimulus-driven magnetization decay (Fig.2g). Our theoretical modeling of baseline and task magnetization decays (Fig. 2g) closely match the observed signal characteristics of the decay and stimulus-evoked HRFs observed in the acquired data (Fig. 2c).

### Comparing in-AB BOLD detection sensitivity between SS and SASS (at 9.4 T)

We investigated the difference in stimulus-evoked BOLD signal sensitivity between a traditional SS paradigm and our SASS paradigm using a simple visual stimulation experiment with awake marmoset monkeys (n=3, Methods) using identical numbers of volumes or stimulus trials. We used an in-AB stimulus presentation so that the whole HRF was captured, even though this is not the optimum for tSNR enhancement. The voxel-based GLM analysis, with a corrected (for multiple comparisons) P < 0.025, revealed activation maps for each individual subject’s runs for both methods, representing stimulus driven activations. At the group level, we observed stronger brain wide activations using the SASS paradigm, compared to the SS paradigm (Fig.3a). SASS showed greater numbers of active voxels at higher z-value compared to those from the SS paradigm. For voxels confined to the primary visual cortex (V1) and medial superior temporal area of cortex (MST), the SASS paradigm showed a similar trend with a greater number of active voxels at higher z-values compared to the SS paradigm (Fig.3b, c). Mean z-score transformed voxel timeseries of each subject run from all active voxels at z-value> 5.1 were used to plot the average timeseries across all subjects, exhibiting characteristic BOLD responses from both paradigms.

**Fig. 3.**
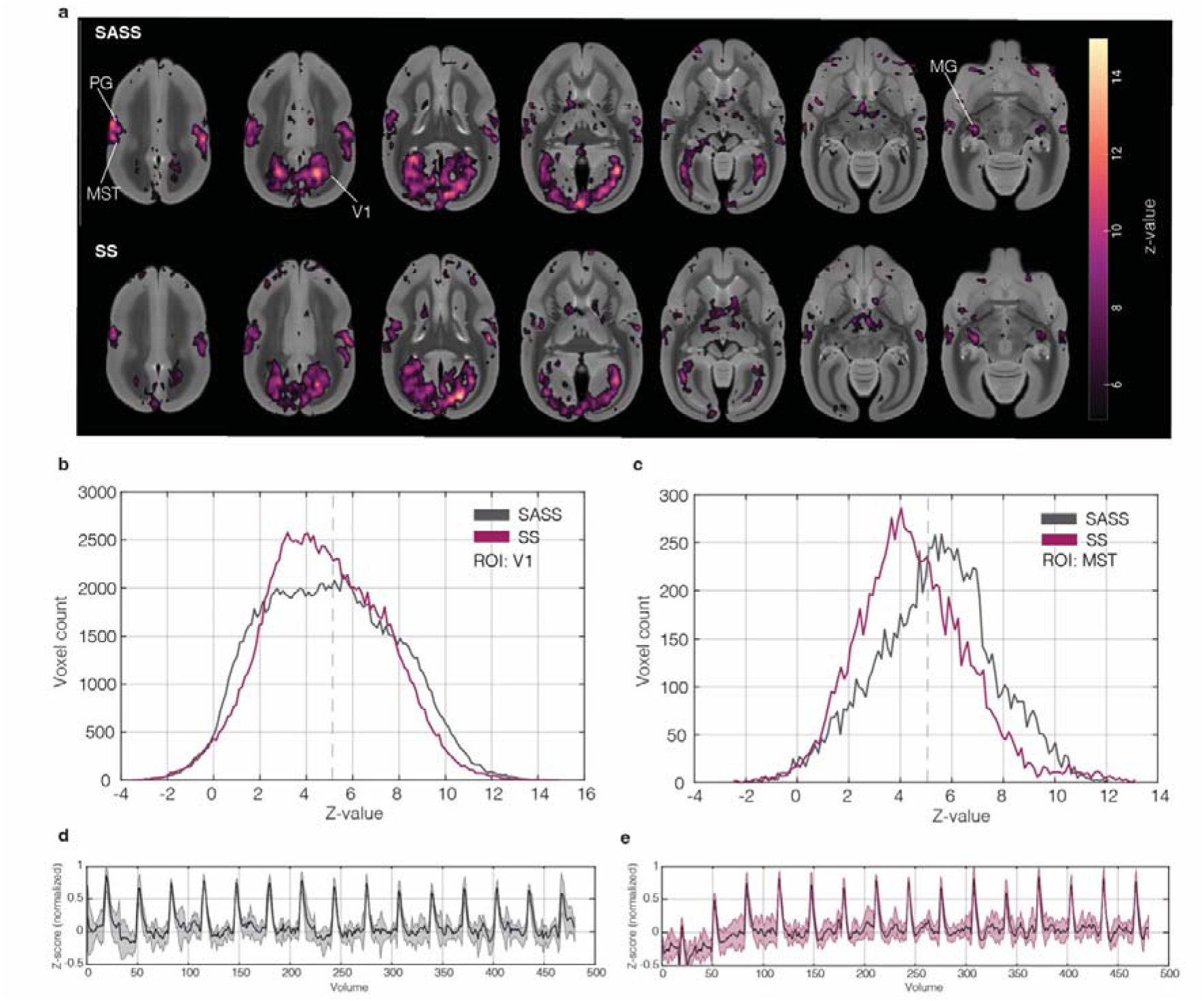
Comparison study with visual stimulus paradigm in marmosets at 9.4 T. **a,** SASS paradigm group activation map (n=3) for an in-AB visual stimulus train and SS paradigm group activation map (n=3, P<0.025, z-value>5.1) from identical visual stimulus paradigm. **b,** Voxel count at respective z-values from group maps of primary visual cortex (V1) for both paradigms, z-value threshold at 5.1 (dashed line). **c,** Voxel count at respective z-values from group maps of medial superior temporal area of cortex (MST) for both paradigms, z-value threshold at 5.1 (dashed line). **d,** Normalized mean z-score timecourse of active voxels above z-value >5.1 (P<0.025) across all subject runs of SASS paradigm (number of volumes per run = 480, TR 1s). **e,** Normalized mean z-score timecourse of active voxels above z-value >5.1 (P<0.025) across all subject runs of SS paradigm (number of volumes per run = 480, TR 1s) (MG-medial geniculate, PG-parietal area).

The mean timeseries show robust responses in the SASS paradigm (Fig.3d) and the SS paradigm (Fig.3e). For both paradigms, the mean z-score timeseries of task regions from active voxels exhibits a characteristic BOLD response across subject runs (Extended Data Fig.2a, b). Based on these observations, we conclude that there exists an enhanced detection sensitivity to BOLD signal changes when utilizing SASS compared to a traditional SS paradigm, when comparing matched numbers of volumes (however, with SASS taking 2 minutes longer than SS due to the AFPs). To match these two approaches by scan time, we also compared 480 volumes with 15 stimulus trials per run for SS paradigm to 352 volumes with 11 stimulus trials per run for the SASS paradigm. Despite having 27% fewer trials, the SASS paradigm still showed better activation maps with higher z-value voxels (Extended Data Fig.1a, b). We extended the decimation of the number of stimulus trials (z-score normalization using predetermined baseline values from 15 baseline ABs) until we achieved a SASS map that was comparable to the full SS map. Only 8 stimulus trials in SASS paradigm were needed to match the activation in V1 ROI from the two paradigms (Extended Data Fig. 1a, b).

### Comparing pre-AB BOLD detection sensitivity between SS and SASS (at 9.4 T)

In our auditory experiment paradigm, high frequency pure tones were presented for 1 second during the MRI noise-free AFPs immediately before the start of each task AB (pre-AB stimulus - Methods). The voxel-based GLM analysis using the truncated stimulus-HRF (Fig.2f), with a corrected P<0.025, revealed z-value maps for each individual run. At the group level (n=2), corrected voxel for a P<0.025 shows robust localized activations in auditory cortex corresponding to the stimulus^10^ (Fig.4a). The mean z-score voxel timeseries from active voxels shows robust BOLD responses to the auditory stimulus presented in the AFP (Fig.4b). Comparison against an auditory stimulus presented in the SS paradigm was not possible due to the superposition of the scanner noise, which leads to very poor maps. To further validate the reliability for detecting a BOLD response to a stimulus presented during the AFP, we replicated our visual stimulus paradigm placing the 1 s stimulus in the AFP (pre-AB stimulus) rather than coincident with the onset of the AB. The resulting group activation map (n=3) (Fig.4c) and time course (Fig.4d) is consistent with our in-AB visual results, noting however that the stimulus conditions are different (pre-AB stimulus presented during a quiet period and in-AB stimulus influenced by scanner noise). For both auditory and visual pre-AB stimulus experiments, the mean z-score timeseries of task regions from active voxels exhibits a characteristic BOLD response across subject runs with an earlier peak (Extended Data Fig.2c, d). From these observations, we conclude that the BOLD response to a stimulus presented during the AFP can be extracted accurately from ABs when an appropriate truncated stimulus-HRF (Fig.2f) is used in the GLM analysis.

**Fig. 4.**
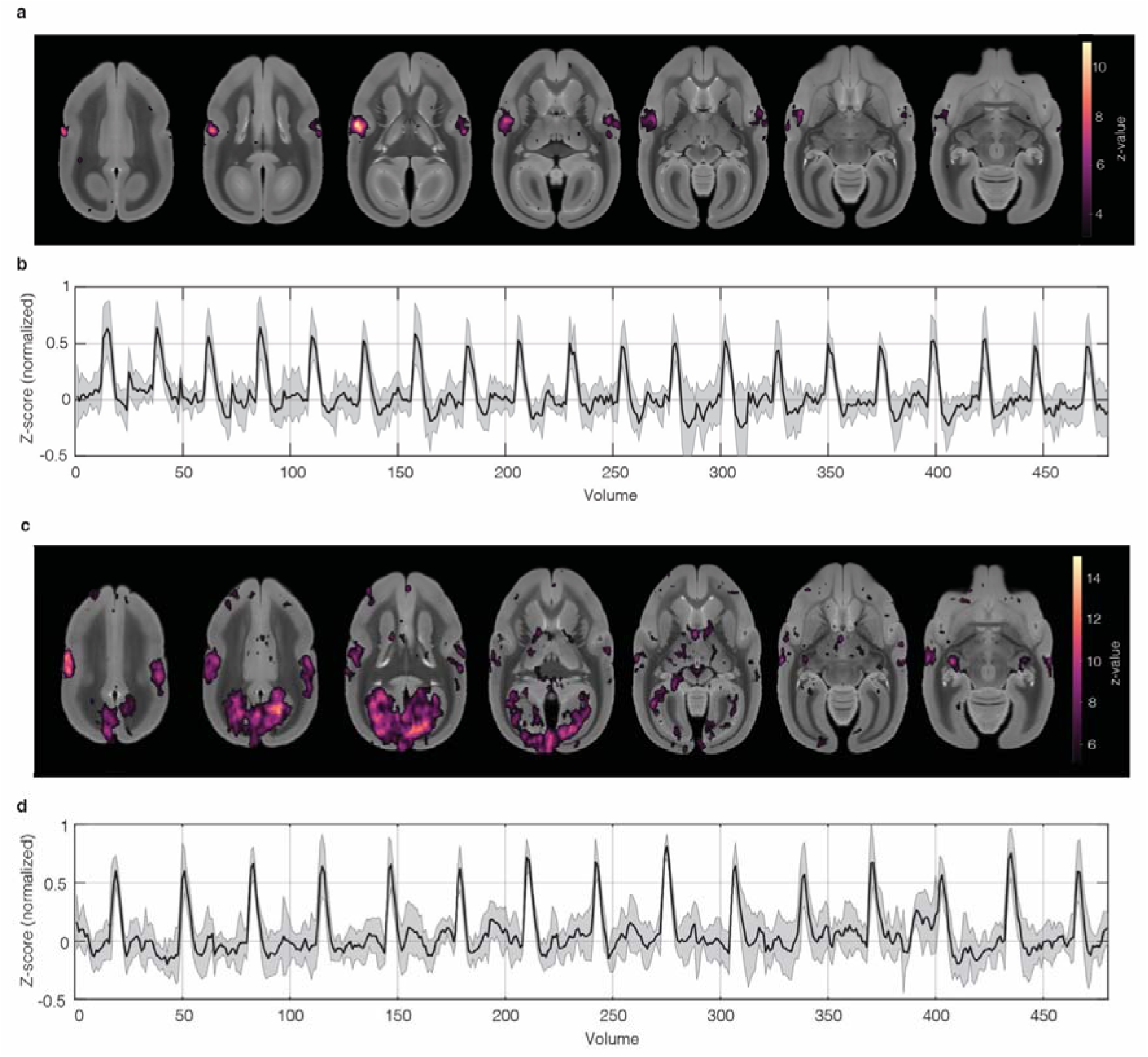
Animal experiments using pre-AB stimulus at 9.4 T. **a,** Group activation maps (n= 2, z-value> 3.1, P< 0 025) for high-frequency pure tone auditory stimuli presented during the AFP using SASS paradigm. **b,** Auditory experiment normalized mean z-score timeseries from active voxels z-value> 3.1 across all subject runs (number of volumes = 480 TR=1 s). **c,** Group activation maps (n= 3, z-value> 5.1, P< 0.025) for pre-AB visual stimuli during silent AFP. **d,** Visual experiment normalized mean zscore timeseries from active voxels above z-value> 5.1 across all subject runs (number of volumes = 480, TR=1 s).

### Comparing pre-AB SASS vs SS in human experiments (at 3 T)

We replicated the SASS and SS paradigms in a 3 T human MRI scanner using a pre-AB presentation of a flickering checkerboard visual stimulus, presented for 1 second in both paradigms (n = 3), with an equal number of volumes and stimulus trials. Cluster-based GLM analysis (corrected *P* < 0.05, *z* > 5.1) revealed robust activation maps in both conditions (Fig. 5a). For the same number of trials (15), the SASS condition produced a greater number of active voxels with higher *z*-values within the visual cortex ROI (Extended Data Fig. 4b), consistent with trends observed in the 9.4 T experiments (Fig.3b, c). To match the scan time of the SS paradigm, we reduced the number of SASS paradigm trials to 11 per run. Despite the reduction, SASS activation maps still showed robust responses with as few as 7 or 11 trials per run (Extended Data Fig. 4a), although not exceeding the SS condition as seen in 9.4 T experiments.

**Fig. 5.**
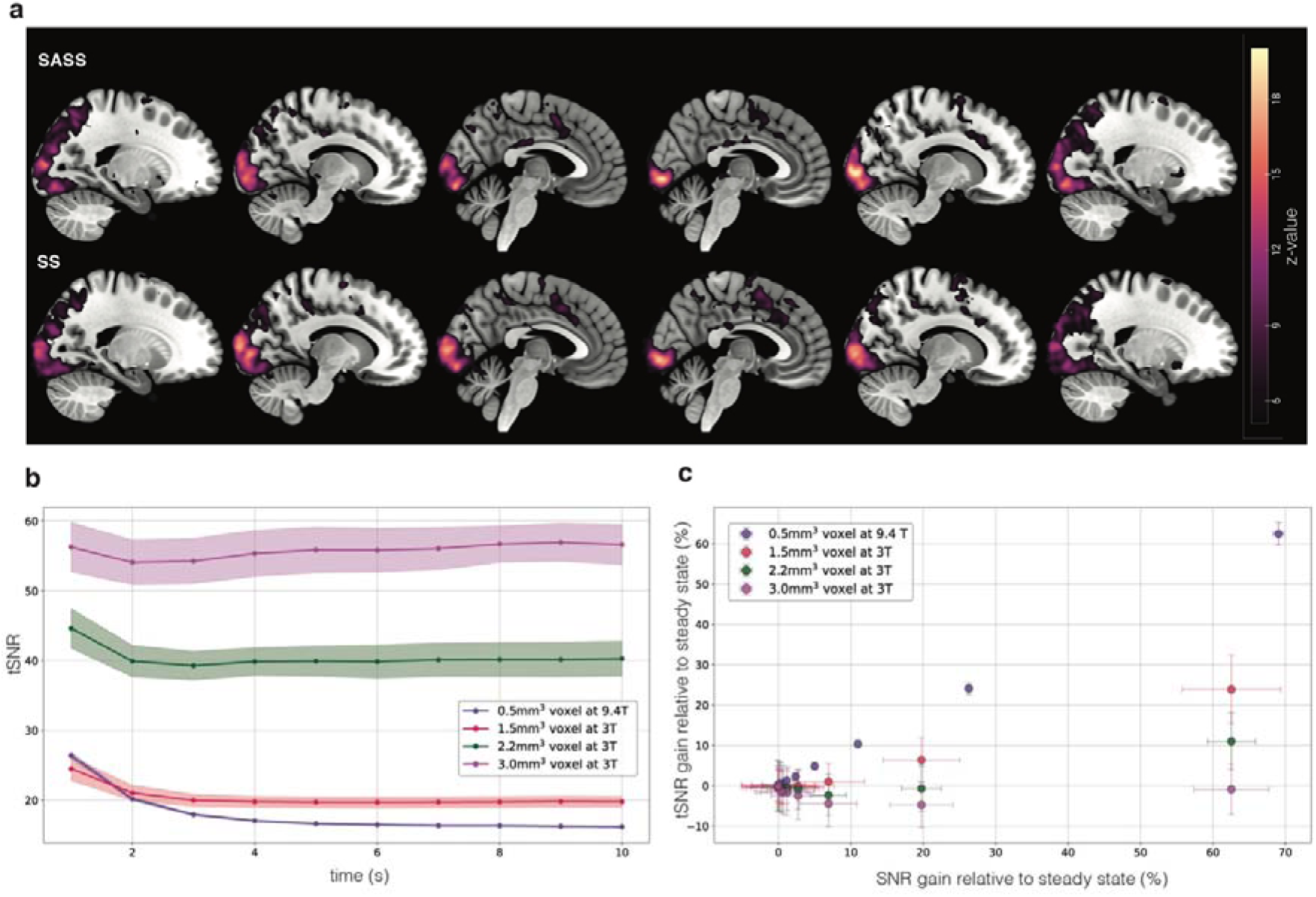
SASS vs SS comparison study for human 3 T and tSNR gains at different voxel resolutions. **a,** SASS and SS z-value maps (n=3, P<0.05, z-value>5.1) from identical flickering checkerboard visual stimulus for 15 stimulus trials per run. **b,** tSNR variation across a 10s long AB (TR =1s) for different voxel resolutions at 3 T (n=3 for all voxel resolutions) includes comparative tSNR values from 9.4 T animal experiment (n=1, 3 runs). **c,** Percent change of tSNR and SNR for each timepoint in an AB compared to their respective steady state values (n= 3 for all voxel resolutions at 3 T, n=1.3 runs for 9.4 T).

### tSNR behavior across noise regimes and voxel resolutions

The benefits of SASS could be highly dependent on whether the tSNR is dominated by thermal noise fluctuations or physiological noise fluctuations. To compare how tSNR varied with image SNR across different noise regimes, we acquired EPI data using the SASS paradigm on a 3 T human MRI scanner at several isotropic voxel resolutions (1.5 mm, 2.2 mm, and 3 mm). The first 10 timepoints of the 9.4 T data were also included in this comparison. Fig. 5b shows the tSNR at each timepoint in the AB for each field strength and voxel resolution (n=3 at each field). The tSNR plots also reveal how the longer T1 relaxation time at 9.4 T vs. 3 T shapes magnetization decay, with the 9.4 T data taking longer to achieve the steady state compared to 3 T. This results in a greater number of timepoints with elevated tSNR above the steady-state value at 9.4 T vs. 3 T.

To assess the dependence of tSNR on SNR, we plotted the tSNR gain relative to steady-state versus the SNR gain relative to steady-state at each timepoint within the AB (Fig. 5c). Plotting the gain relative to the steady-state values (mean of last 3 points in AB) avoided the complication of making absolute SNR measurements that need to be corrected for coil g-factors, preamplifier gain and Rician noise. Normalized magnetization decay profiles are expected to be consistent across 3 T voxel resolutions due to identical FA and TR values. However, minor deviations occur at larger voxel sizes, likely due to partial volume effects (Extended Data Fig. 5). To correct for relative SNR differences, normalization of the intensity relative to the 1.5 mm voxel was applied at each timepoint. This adjustment compensates for increased CSF and WM contamination in larger voxels, with 1.5 mm voxels exhibiting the least partial-volume influence. Fig. 5c highlights the relative tSNR gain versus the relative SNR gain for each resolution. At 9.4 T with 0.5 mm isotropic resolution, tSNR increases linearly with SNR with near unity slope (0.82, R^2^ = 1.0). At 3 T the tSNR dependence on SNR progressively decreases as voxel size increases, consistent with moving from a thermal to physiological noise dominated regime. At 3 mm resolution, the tSNR is independent of the SNR, indicating a physiologically dominated noise mechanism.

## DISCUSSION

### SASS design considerations

MRI noise-free regions in periodic sequences operating in the steady-state regime have been previously utilized to facilitate auditory stimulus presentation^11–14^. Such approaches use either sparse sampling or continue applying the radio frequency (RF) pulses to maintain the steady-state of the magnetization during the quiet periods. Sparse sampling limits the temporal resolution of the HRF, while the RF in the silent periods is in fact difficult to implement for the ubiquitous slice-multiplexed fMRI pulse sequences^15,16^ such as multi-band excitations which require gradients^17^ and interfere with multi-modal fMRI, such as fMRI-EEG. Our approach is distinct in that we use the AFPs to both fully recover the magnetization and present stimuli, after which we read out the fully-relaxed BOLD response to the stimulus with an AB. This approach compensates for the reduced time efficiency of incorporating AFPs by exploiting the enhanced tSNR during the approach to steady-state. At a minimum, the AFP just needs to be long enough to allow full recovery of the magnetization, namely at least 4 times the T1 of the grey matter. The stimulus should be placed such that the peak of the BOLD response occurs during the period of enhanced tSNR, i.e. the initial part of the AB. The earlier the BOLD signal peaks during the magnetization decay, the higher the peak detection sensitivity. This places some constraints on the duration and location of the stimulus in the SASS paradigm design. A short duration (few seconds) stimulus placed in the AFP immediately before the start of the AB is appropriate. This type of short stimulus presentation paradigms is characteristic of event-related designs in fMRI studies. We also note that the time-to-peak for the BOLD response is shorter in smaller species. This means the stimulus can be placed further away from the start of the AB for human experiments, allowing slightly longer presentation durations.

### SASS signal enhancement

Through judicious presentation of the stimulus in the AFP, the peak of the BOLD HRF can be timed to be in the AB region where tSNR at the first point is up to ∼60% higher than the steady-state tSNR (at 9.4 T). Integrating the volume-by-volume tSNR values multiplied by the volume-by-volume HRF yields a weighted tSNR gain of 22% to 36%. Such an increase in tSNR should yield an approximately doubling of detection efficiency for our chosen parameters at 9.4 T, as we observed (Extended Data Fig. 1). At 3 T the efficiency gains are reduced, with 11 SASS trials producing equivalent maps to 15 SS trials (Compare Fig. 5a with Extended Data Fig. 4), consistent with the tSNR increase of ∼12% ± 4%. This reduction occurs because the number of points on the decay curve of the tSNR at 3 T that are above the SS magnetization value are fewer due to the shorter T1 (Fig. 5b) even though the magnetization decays are quite similar (Extended Data Fig. 5). Fig. 5c shows that 5 points are above the SS tSNR baseline at 9.4 T, but only 2 (2.2 mm, 3 mm) to 3 (1.5 mm) are above this baseline at 3 T.

### Noise estimation for tSNR

In the demonstrations here we have used equal numbers of alternating task and baseline ABs. This is not a requirement of the SASS paradigm. One simply needs enough baseline ABs to accurately determine the temporal standard deviation of each volume in the AB to allow for accurate z-score normalization, either acquiring them in between task ABs (as done here) or having an independent run to characterize the baseline AB and then using only task ABs in subsequent runs. However, to simplify (in a statistical sense) the z-scoring process, the number of volumes in the task and baseline ABs should ideally be the same. The tSNR in the steady-state is the same for every volume, therefore only point in that equilibrium is needed to z-score all the images acquired in the steady-state. In our experiments, we used a longer than required AB to clearly demonstrate the SASS gains, but the AB can be made shorter, governed by the HRF characteristics. Acquisition of baseline ABs can be completely forgone if other approaches to estimating the tSNR in a voxel are used, such as those used in rapid event-related fMRI. We did not use those approaches here to allow precise comparison between SS and SASS.

### Pre-AB vs in-AB sensitivity differences

We note the almost doubling of the peak detection sensitivity of the pre-AB visual stimulus with the in-AB visual stimulus at 9.4 T. These are not quite the same experiment from a psychophysical perspective, the major difference being that in the pre-AB condition the visual stimulus is presented in a period of silence before the AB starts. The presentation of temporally congruent auditory and visual stimuli in the in-AB condition, is known to enhance activation in the sensory cortices and elsewhere due to multimodal integration, inhibition and disinhibition^18–20^. These arguments should make for a more robust in-AB map, yet the pre-AB map is visually comparable or superior in part because of the presentation of the visual stimulus in isolation and in large part because the HRF peak of such a pre-AB stimulus plays out during the highest tSNR section of the magnetization decay, while the in-AB HRF occurs at lower tSNRs.

### Noise regime

With the RF coil and acquisition parameters employed in the 9.4 T experiments, the imaging operates firmly within the thermal noise regime, where tSNR scales linearly with available magnetization (i.e. SNR). This regime is characteristic of many ultra-high-field laminar fMRI studies conducted in both humans and animals. Notably, in human studies at 7 T, the T1 relaxation time of grey matter^21^ (∼2 seconds) is comparable to that of marmoset grey matter at 9.4 T, which would result in similar magnetization recovery dynamics and tSNR gains to those observed in our animal study. In contrast, at 3 T or coarser spatial resolutions, voxels enter a mixed or physiological noise regime, where tSNR depends progressively less on the SNR as the voxel size increase^22^. The results show the highest gain in tSNR for smaller voxels, where we expect lower physiological noise influence compared to the larger voxels. At the 2.2 mm isotropic resolution used in our SASS human visual experiments (typical of modern 3 T fMRI acquisitions), we still observe tSNR gains while achieving whole-brain coverage with a 1-second TR.

Together, Fig. 5b and 5c illustrate the distinct relationships between tSNR and magnetization that emerge across noise regimes, consistent with previously established models. These findings underscore the influence of voxel size and underlying noise composition in determining the effectiveness of the SASS paradigm. We note that for an equivalent number of trials, under no conditions did the SASS paradigm underperform a conventional SS paradigm. In the worst case (3 T, 3 mm) the two approaches were equivalent.

These gains could be further enhanced by synchronizing stimulus presentation/data acquisition with a physiological signal to reduce the physiological noise contribution. The ability to synchronize the stimulus presentation and subsequent AB to a parameter such as cardiac and/or respiration phase could increase the tSNR in fMRI experiments^23^. Likewise, triggering acquisitions based on a behavioral event (e.g. eye fixation, saccade, key press or EEG waveform) could significantly decrease inter-trial variability. This approach would enable image acquisition to occur solely in response to triggers from the subject, potentially giving the subject control over the stimulus presentation.

### Benefits and limitations of an AFP

The imaging gradients and the radio frequency pulses are not present during the AFP. This has many advantages if the stimulus is presented in the AFP. Auditory signals or instructions can be presented to the subject with no competing MRI sound interference. Speech or vocalization responses can be recorded with high fidelity during this time as well. In multi-modal experiments, such as simultaneous electrophysiology and fMRI^24–26^, electrophysiology data acquired are heavily affected by MRI gradient artifacts and it remains challenging to clean the data and maintain diagnostic waveform fidelity. By presenting a stimulus in the AFP, we enable the unique ability to record undistorted single-unit, local field potential or evoked potential responses to the stimulus without the interference of MR gradient or RF artifacts. The AB following the AFP can still capture the BOLD response, because the response peaks with a hemodynamic delay of several seconds. Alternatively, where the stimulus and response are far apart, such as in a working memory task, the fMRI acquisition can be triggered by the stimulus or the response. Incorporating stimulus presentation during the AFP offers other additional benefits, such as minimizing stimulus-driven motion artifacts in experiments where immediate motion due to the stimulus (e.g. a button press or joystick movement) does not affect the ABs that follow. Because the sequence can be started at will after a sufficiently long AFP, it lends itself to fMRI studies of neonates and pediatric subjects, where it can be triggered when the subject is not moving while maintaining silence the rest of the time.

Since the baseline is derived from non-task ABs, there is no real need to acquire an AB longer than the number of volumes where the tSNR is higher than the steady state, i.e. roughly 5-8 seconds. The peak and width of the BOLD response to a short stimulus is well matched in time to the decay of the magnetization towards steady-state in the AB. The actual stimulus-induced HRF has ample time to recover during the AFP.

Multiple repetitions of the imaging volume with a short enough TR to well characterize the BOLD response within an AB can be challenging, depending on the required spatial resolution and coverage. We show that using modern multi-band fMRI approaches, it is certainly possible to obtain whole brain fMRI volumes in ∼1 s at cortex-respecting spatial resolutions on human MRI systems at 3 T^16,27^. However, whole brain imaging at millimeter or submillimeter resolution at 7 T isn’t feasible with current scanners and software and coverage or temporal resolution would have to be compromised. Alternately, the stimulus can be temporally jittered between presentations to effectively subsample the BOLD response. SASS can be used for long stimulus durations in the AFP, however, it can only capture the final few seconds of the stimulus response.

The use of the SASS paradigm has could benefit enhanced tissue discrimination for grey matter segmentation directly from the functional images, without the reliance of a T1 weighted image. This is because tissue components such as grey matter and white matter decay with different T1 relaxation time constants and can therefore be segmented from their decay curve properties. By using the tissue segmented and higher contrast from SASS images, one could improve the challenging registration between the usually distorted functional and higher-fidelity structural images. The segmentation could be further improved using timeseries clustering or curve fitting methods, possibly allowing for differentiating not just grey or white matter voxels but also larger blood vessels^7^. We note however that at lower voxel resolutions, this delineation is not accurate due to partial volume effects from GM, WM and CSF in the voxel yielding multiple, mixed magnetization components.

In general, thermal noise dominated event-related fMRI experiments are expected to benefit most strongly from the SASS approach. In the physiological noise regime, SASS at least matches the sensitivity of an SS acquisition, while still allowing the other benefits for an AFP described above. SASS may be particularly helpful for signal-starved human 7 T layer fMRI studies, where the tSNR gain in acquired volumes becomes particularly valuable in delineating layer differences in BOLD activation.

## METHODS

### Animal preparations

Experimental procedures followed the guidelines outlined by the Canadian Council of Animal Care policy and were conducted under a protocol approved by the Animal Care Committee of the University of Western Ontario #2021-111. We ensured all ethical regulations governing animal use are followed. Ultra-high field fMRI data used for all experiments were obtained from 6 awake common marmoset monkeys (*Callithrix jacchus*): two females and four males. (6.7±3.7 years and 403±33g) (Visual experiment n = 3, auditory experiment n = 2, tSNR calculation n = 1).

To prevent head motion during awake MRI acquisition, animals were surgically implanted with an MR-compatible machined PEEK (polyetheretherketone) head post^28^, conducted under anesthesia and aseptic conditions. During the surgical procedure, the animals were first sedated and intubated to ensure they remained under gas anesthesia, maintained by a mixture of O2 and isoflurane (0.5–3%). With their heads immobilized in a stereotactic apparatus, the head post was positioned on the skull following a midline skin incision along the skull. The head post was secured in place using a resin composite (Core-Flo DC Lite; Bisco). Heart rate, oxygen saturation, and body temperature were continuously monitored throughout the surgery. Two weeks post-surgery, the monkeys were acclimated to the head-fixation system and the MRI environment through a three-week training period in a mock scanner^29^.

### 9.4 T MRI and fMRI procedures

MRI experiments were performed on a 9.4-T/31-cm magnet (Agilent, Palo Alto, CA, USA) interfaced to a Bruker Avance NEO console (Bruker BioSpin Corp, Billerica, MA) and equipped with an in-house designed and built 15-cm gradient coil of 450mT/m strength^30^, with Paravision-360 v3.3 software at the Centre for Functional and Metabolic Mapping located within the Robarts Research Institute at the University of Western Ontario. An 8-channel phased-array receive coil was used in combination with a birdcage transmit coil for data acquisition.We employed a gradient-echo-based single-shot echo-planar imaging (EPI) sequence with following parameters: repetition time (TR) = 1s, echo time (TE)= 15□ms, flip angle = 40°, field of view (FOV) = 48×64 mm^2, matrix size = 96×128, isotropic resolution of 0.5□mm^3^, 25 coronal slices, bandwidth = 400□kHz, and a GRAPPA acceleration factor of 2 (left-right). Additionally, an extra set of EPIs with opposite phase-encoding direction (right-left) were acquired for EPI-distortion correction purposes. For each session, the T2-weighted anatomical images were acquired for each session using the TurboRARE2D pulse sequence (6 averages, 42 slices, slice thickness = 500 μm, FOV 48×64 mm^2, matrix size 192×192, TE = 39 ms, TR = 7 s, echo spacing = 13 ms, and RARE factor 8).

For the SASS paradigm built from the above-mentioned EPI sequence, we incorporated recurring 8-second-long AFPs after every 16 (visual stimulus paradigm) or 12 (auditory stimulus paradigm) second long ABs. The inclusion of AFPs increased the scan time compared to a SS paradigm as the number of volumes acquired was kept the same in both sequences (480 volumes in each run). For scan time comparison with SS paradigm, we reduced the number of volumes per run in the SASS paradigm to best match the duration of the SS paradigm. The number of slices in both visual and auditory paradigms were set to 25 slices, covering respective regions of interest according to the stimulus paradigm. For the visual stimulus paradigm, data from each animal was collected during a single session in the given order: SASS paradigm (in-AB stimulus), SS paradigm, SASS paradigm (pre-AB stimulus).

### MRI and fMRI procedures (human experiments)

Human experiments were conducted following the Western University Health Science Research Ethics Board (HSREB) approved ethics (REB:124962) and after obtaining signed consent from all participants. MRI data were acquired on a 3 T Siemens MAGNETOM Prisma Fit scanner (Siemens Healthineers, Erlangen, Germany) located at the Centre for Functional and Metabolic Mapping located within the Robarts Research Institute at the University of Western Ontario. The scanner was equipped with a 32-channel head coil and operated using the Syngo MR XA60 software. High-resolution T2-weighted anatomical images were acquired using a 3D MPRAGE sequence with 1 mm isotropic resolution (TR = 2.3s, TE = 2.98ms, TI = 900ms, flip angle = 9°). GRAPPA acceleration x2 was used. The acquisition provided full-brain coverage with 256 × 256 matrix size. Images were acquired in the sagittal plane. Whole-brain blood oxygenation level–dependent (BOLD) images were collected using a modified (from the product) 2D simultaneous multi-slice (SMS) EPI sequence with the following parameters: repetition time (TR) = 1000ms, echo time (TE) = 30ms, flip angle = 62°, SMS acceleration factor = 3, pixel bandwidth = 1860 Hz/pixel, GRAPPA acceleration factor = 3 and partial Fourier = 6/8. Each volume comprised 48 axial oblique slices acquired in an interleaved order, with isotropic voxel size = 2.2 mm³. and an in-plane FOV = 211 mm covering the brain. The phase encoding direction was posterior–anterior (denoted j− in JSON files). Additionally, an extra set of EPIs with opposite phase-encoding direction (denoted j in JSON files) were acquired for EPI-distortion correction purposes. For tSNR calculations, we conducted another experiment using the same modified SMS EPI sequence to acquire volumes at different voxel resolutions. The data were collected at 1.5 x 1.5 x 1.5 mm (pixel bandwidth= 1984Hz/pixel, 36 slices), 2.2 x 2.2 x 2.2 (pixel bandwidth= 1860 Hz/pixel, 48 slices) and 3 x 3 x 3 mm (pixel bandwidth= 1880 Hz/pixel, 48 slices) resolutions.

For the visual SASS paradigm (n=3), the EPI sequence described above was used with recurring 8-second-long AFPs following each 16-second-long AB. The same sequence, excluding the AFPs, was used for the visual SS paradigm. To maintain consistency across paradigms, the number of acquired volumes and the number of stimulus trials were held constant (15 stimulus trials, 480 volumes per run), despite the increased scan time in the SASS paradigm due to the inclusion of AFPs. Two runs per subject for either SASS or SS paradigms were acquired in a single session. For tSNR analysis (n=3), data were collected using the EPI sequence with recurring 8-second-long AFPs following each 10-second-long AB. Resting-state data were acquired using this configuration, with 500 volumes per run (comprising 50 ABs of 10 volumes each).

### Acquisition blocks and acquisition free period

We introduced Acquisition-Free Periods (AFPs) to induce magnetization relaxation within an fMRI run. AFPs represent silent periods in the sequence where no volumes are acquired, and no RF is applied. The duration of the AFP is made long enough for full magnetization recovery, typically 4-5 times the T1 relaxation time constant of grey matter (1.3-1.6s in humans at 3 T and 2.0-2.2s in marmosets at 9.4 T)^31,32^. The volume acquired immediately after an AFP captures the magnetization at its maximum, with the subsequent volumes capturing the magnetization decay before reaching a steady-state intensity governed by the T1 and FA. The acquisition period following the AFP is called an acquisition block (AB), within which a series of volumes are acquired with a specified repetition time (TR) and FA. ABs, each of identical length, form independent periods of the time series which serve as the basic repeating unit for designing task paradigms. The AFP and AB durations can be modified according to the task design making sure there is enough time for full magnetization recovery and characterization of the BOLD hemodynamic response (only if necessary).

For all our fMRI experiments, in both marmosets and humans, ABs are categorized into alternating task and baseline blocks. The length of ABs is tailored to capture the hemodynamic response associated with a stimulus presented either before (pre-AB stimulus) or within a task block (in-AB stimulus). We use baseline blocks to characterize the magnetization decay curve, permitting the z-score normalization of task blocks for subsequent analysis. For our respective stimulus paradigms, we employed two different block lengths (16 seconds for visual experiment (humans and marmosets) and 12 seconds for auditory experiments (marmosets)) with the same minimum AFP duration of 8 seconds (∼4*T1).

### Theoretical modeling for 9.4 T

The relaxation process of the magnetization vector in the presence of an external magnetic field is described by the Bloch equations^6^. In the context of T2*-weighted EPI acquisitions, our solutions derived from the Bloch equations for the magnetization *M*_(*n*)_ after the *n*^th^ excitation in a series of excitations is governed by imaging data for the magnetization relaxation values at each *TR* (1 second) for an AB length of 16 and tissue parameters given in the equation (Equation 1). Using the equation, we simulated fully relaxed condition) value was set to 100. We used a *θ* (flip angle) of 50 degrees (Ernst seconds (Fig.2c). The maximum magnetization *M*_0_ (after a single 90-degree pulse from the angle) in the simulation to match the nominal FA of the experiment over the slice. The grey matter *T*_1_ time was set to 2.2 seconds matching the grey matter *T*_1_ relaxation time constant in marmosets ranging between 2.0 and 2.2 seconds at 9.4 T MRI.

To simulate the influence of a stimulus on the decay, we generated an HRF using the double gamma function defined in the FSL toolbox (time to peak - 3 s and time to peak of undershoot - 8 s). The maximum BOLD percent signal change due to the stimulus was fixed to 4 % (typical signal change observed in 9.4 T fMRI experiments). The baseline magnetization curve was multiplied with this HRF function convolved with a stimulus duration of 1 second. The resultant magnetization values represent a stimulus-modulated approach to steady-state. Because of this multiplication, the initial magnetization values during a task are slightly higher than those during the baseline as the BOLD signal is a few percent higher than baseline (Fig.2g). When the stimulus occurs during the AFP, the generated HRF should be truncated before multiplying with the baseline magnetization, as the onset of stimulus doesn’t coincide with the start of AB (Fig.2f).

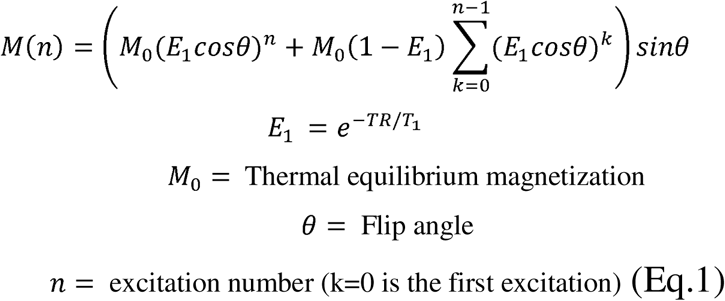

### Temporal Signal-to-Noise (tSNR) calculation (animal experiment)

tSNR values in AB volumes were calculated using preprocessed data from a single subject across four runs (30 ABs per run). For each timepoint in an AB, the mean and standard deviation were computed for each voxel across all blocks. Independent Component Analysis (ICA) using FSL’s melodic ICA was applied to the time series, revealing inherent differences in magnetization decay between grey and white matter voxels (Extended Data Fig.3a). This resulted in independent components corresponding to grey and white matter voxels (Extended Data Fig.3b). The white matter components were removed, and the mean grey matter temporal signal-to-noise ratio (tSNR) was calculated using the mean and standard deviation obtained for each timepoint in an AB (Fig.1c).

### tSNR gains across noise regimes

Temporal signal-to-noise ratio (tSNR) was calculated using motion-corrected functional MRI data (n=3). Each run consisted of 50 ABs (500 volumes) with each AB spanning 10 seconds. Data were collected at three isometric voxel resolutions: 1.5 mm, 2.2 mm, and 3 mm. For each timepoint within an AB, the mean and standard deviation of the BOLD signal were computed across all ABs on a voxel-wise basis. Grey matter voxels were segmented from each subject’s T2-weighted anatomical image using FSL’s FAST segmentation tool. Functional images were registered to the anatomical space, and tSNR values were computed within the segmented grey matter mask for each voxel resolution.

tSNR variation across AB volumes was calculated for each voxel resolution (Fig. 5b), including comparative data from 0.5 mm isotropic resolution at 9.4 T (animal experiments). To assess the relationship between tSNR and signal magnitude, we computed the percent change in tSNR and signal intensity at each timepoint within an AB, relative to the steady-state value (mean of the final three timepoints).To adjust for the minor deviations in intensity occurring at larger voxel sizes, likely due to partial volume effects, intensity ratios between the 1.5 mm and larger voxels were multiplied at each timepoint in the AB. Fig. 5c shows the percent change in magnetization plotted against tSNR across resolutions, highlighting distinct SNR–tSNR relationships under different noise regimes.

### Baseline z-score normalization of acquisition blocks

To extract the BOLD signal changes evoked by the stimulus, we needed to eliminate the influence of the T1 governed magnetization decay from the ABs. Prior methods have utilized a global decay regressor for regressing the decay^33^, which is inappropriate due to the inherent variability in T1 decay curves across different voxels, influenced by their tissue characteristics and RF coil inhomogeneity that causes variable FAs. For SASS paradigm runs, we utilized alternating baseline blocks within each run to standardize all blocks uniformly. Within each run, standard preprocessed fMRI volumes are initially segmented into task and baseline blocks. For each voxel timepoint within the AB, the mean and standard deviation are calculated from all baseline blocks. These statistical measures are then utilized to calculate standardized z-scores for both individual task and baseline ABs (Equation 2). This transformation enables the expression of voxel signal fluctuations between each time points in terms of their respective standard deviation units from the baseline mean values, thereby eliminating the decay influence from each block timepoint. For each voxel the resultant z-scores are mean shifted to their respective temporal mean from all baseline blocks. Adding the mean as a constant was only necessary to be able to utilize the conventional FSL tools (for registration of subject’s functional to anatomical to template space). All z-score transformed blocks are concatenated accordingly and are fit for further analysis using the FSL toolbox. The same method was used to z-score normalize SS paradigm data by utilizing the well-defined baseline and task regions in the raw timeseries. For SS runs, the first baseline region (16 volumes) during approach to steady-state was replaced with the subsequent baseline region before normalization.

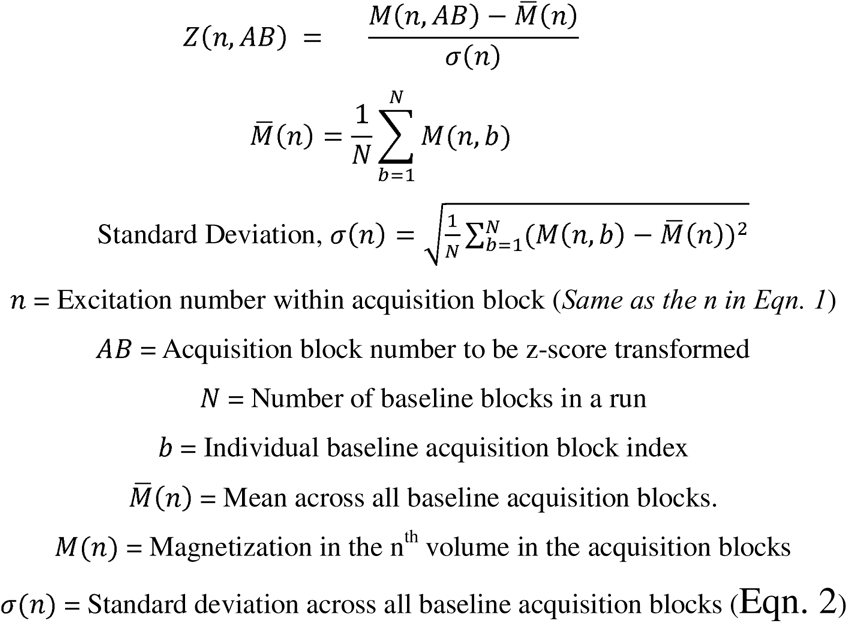

### Stimulus design for animal experiments

We utilized two different stimulus designs for the study. For all visual stimulus experiments, a 10 Hz flickering LED light was presented for a duration of 1 second. Fifteen task ABs were collected in each run, interleaved with 15 baseline ABs. The ABs were 16 s long. For the SASS paradigm, two different stimulus designs were used such that the stimulus was presented coincident with the start of the AB (in-AB stimulus) or during the AFP immediately before task AB (i.e. end of stimulus coincident with the start of task AB) for the other (pre-AB stimulus). For the visual experiment with the SS paradigm, the stimulus was presented every 32 seconds allowing well-defined baseline and task regions (16 s). The FA and the TR was the same in both SASS and SS runs so that the steady-state magnetization was identical.

For the auditory experiments using SASS paradigm, we utilized an AB length of 12 seconds with auditory stimulus presented during AFP (pre-AB stimulus). The design had alternating baseline and task ABs, with a total of 20 stimulus trials in each run. We presented alternating (50ms ON/OFF) high frequency pure tones (16 kHz – 20 kHz) for 1 second every 40 seconds before the start of each task AB, which includes the 8 s long AFPs between ABs.

The monkeys were positioned in a horizontal MR system operating at 9.4 T, adopting a sphinx position within an MRI-compatible restraint system with their heads secured using a head post^29^. An MR-compatible camera (model 12M-i, MRC Systems GmbH, Heidelberg, Germany) was integrated in the MRI chair to monitor the animals during data acquisition. Both the onset of visual and auditory stimuli was synchronized with the MRI using TTL (transistor-transistor logic) pulses from the scanner and a custom-written Python program running on a Raspberry Pi 4 model B (Raspberry Pi Foundation, Cambridge, UK). The TTL pulses were utilized to ensure precise synchronization of stimulus onsets and volume acquisition. For the visual stimuli, we custom-built a setup with two bilaterally attached white LEDs, positioned approximately 10 cm from the animals’ head location. For the auditory stimuli, following the head fixation steps, MRI-compatible auditory tubes (S14, Sensimetrics, Gloucester, MA) were inserted into the animals’ ear canals bilaterally and secured using sound-attenuating silicone earplugs (Amazon) and self-adhesive veterinary bandage^29,34^.

### Data analysis and statistics for animal experiments

The DICOM functional images were converted to NifTi format using AFNI’s^35^ dcm2niix function. EPIs acquired with opposite phase-encoding direction were used for EPI distortion correction (top-up) using custom-written MATLAB code. Using the FSL toolbox^36^, the raw functional images were preprocessed by applying SUSAN noise reduction, MCFLIRT motion correction and spatial smoothing at full width at half-maximum Gaussian kernel (FWHM) of 0.65 mm. For each run, brain extracted functional images were linearly registered to the respective manually skull stripped T2-weighted anatomical images of the animal using FSL’s FLIRT function. The T2-weighted anatomical images were registered to the NIH marmoset brain atlas^37^ via affine linear registration (12 degrees of freedom) using FSL’s FLIRT function. All preprocessed fMRI data for both SS and SASS paradigms were then z-score normalized using the baseline ABs or regions within each run and mean shifted to respective baseline temporal mean values.

The z-score transformed fMRI runs were used for statistical analysis using the general linear model (GLM) approach in the FSL toolbox^36^. The data underwent slice-time correction as the slices were acquired in interleaved order (0, 2, 4, …1, 3, 5, …). For comparison study between SASS (in-AB) and SS paradigms, we used a gamma HRF peaking at 3 seconds (with temporal derivates to find the best fit) for generating a stimulus onset regressor for GLM analysis. For our pre-AB stimulus condition, where the stimulus is presented immediately before the start of task AB, we use a custom truncated stimulus-HRF (without time-derivatives) (Fig.2f) to match the stimulus onset to capture the characteristic BOLD peaking a second earlier than in-AB stimulus. The pre-AB analysis used a fixed HRF (shown in Fig.2f) due to limitations in the analysis software package for incorporating time-derivate for custom HRFs. For each individual run, corrected voxels (Gaussian Random Field Theory (GRF) correction for multiple comparisons using one-tailed t-test) with a 5.1 (visual) and 3.1 (auditory) z-value thresholds at P<0.025 were considered active voxels. The z-value maps were generated per run for each subject for all different experimental conditions. These maps were used for FSL’s fixed-effects group analysis for each condition resulting in group z-value maps for P<0.025. All the group z-value maps were registered to the NIH marmoset brain atlas^37^ template space to visualize the differences in voxel responses across difference conditions.

### Data analysis and statistics for human experiments

DICOM functional images were converted to NifTi format using AFNI’s^35^ dcm2niix function. EPIs acquired with opposite phase-encoding direction were used for EPI distortion correction (top-up) using custom-written MATLAB code. Using the FSL toolbox^36^, the raw functional images were preprocessed by applying SUSAN noise reduction, MCFLIRT motion correction and spatial smoothing at full width at half-maximum Gaussian kernel (FWHM) of 4 mm. For each run, brain extracted functional images were non-linearly registered to the respective skull stripped T2-weighted anatomical images of the animal using FSL’s FLIRT function. The T2-weighted anatomical images were non-linearly registered to the FSL’s human MNI152 template using FSL’s FLIRT function. All preprocessed fMRI data for both SS and SASS paradigms were then z-score normalized using the baseline ABs or regions within each run and mean shifted to respective baseline temporal mean values (to retain brain structures for registration purposes).

The z-score transformed fMRI runs were used for statistical analysis using the general linear model (GLM) approach in the FSL toolbox^36^. For the comparison study between SASS (pre-AB) and SS paradigms, we used a gamma HRF peaking at 6 seconds for generating a stimulus onset regressor for GLM analysis. For our pre-AB stimulus condition, where the stimulus is presented 4 seconds before the start of each task AB, we use a custom truncated stimulus-HRF to match the stimulus onset to capture the characteristic BOLD peak. The pre-AB analysis used a fixed HRF due to limitations in the analysis software package for incorporating a time-derivate for custom HRFs. For each individual run, corrected voxels (Gaussian Random Field Theory (GRF) correction for multiple comparisons using one-tailed t-test) with 5.1 z-value thresholds at P<0.05 were considered active voxels. The z-value maps were generated per run for each subject for all visual experimental conditions. These maps were used for FSL’s fixed-effects group analysis for each condition resulting in group z-value maps for P<0.05. All the group z-value maps were registered to the human MNI152 template (1 mm^3^) space to visualize the differences in voxel responses across difference conditions.

### Stimulus design for human experiments

For visual stimulus experiments in human subjects, an 8□Hz flickering checkerboard (8×8 squares) was presented for 1□second. Each run included 15 task ABs interleaved with 15 baseline ABs, with each AB lasting 16□seconds. Each AB was followed by an 8-second-long AFP. In the SASS paradigm, the stimulus was presented 4□seconds prior to the onset of each task AB (during AFP), triggered by TTL pulses from the scanner. In the SS paradigm, the stimulus was presented every 32□seconds, enabling similar well-defined baseline and task periods (16□s). The scanning parameters were identical across SASS and SS runs to ensure consistent steady-state magnetization. Custom Python scripts were used to synchronize stimulus presentation with TTL pulses from the scanner. Participants lay supine in the MRI scanner and viewed stimuli via a rear-projection system (Avotech SV-6011) reflected through a mirror mounted on the head coil.

## Data availability

The raw MR data acquired, and sequence protocol used for the study is available from the corresponding author upon reasonable request. Example data raw and filtered functional Nifti files are available for download at Open Science Forum (OSF) repository.

## Code availability

Custom python and web-based generator for stimulus paradigm, simulation and z-score normalization are available at Open Science Forum (OSF) repository (https://osf.io/p9x8k/?view_only=14fc69dd900144b0a54ed69603c80179).

### Acknowledgements

We would like to thank all the members of the Centre for Functional and Metabolic Mapping (CFMM) at Robarts Research Institute, Western University for their valuable contributions to this project. We thank Dr Alex Li for valuable input on sequence design and acquisition protocol. We thank Dr Kyle Gilbert and Peter Zeeman for the design and building RF coil, MR animal chair and stimulus paradigm setup. We thank our veterinary staff Miranda Bellyou, Whitney, Cheryl and Hannah for their help with conducting the animal experiments. I would like to thank Joe Gati and Dr Atena Akbari for their support in setting up human experiments. I would like to show my gratitude to all the participants the 3 T MRI technicians Oksana Opalevych, David Reese and Trevor Szekeres who helped acquire the data. This work was supported by funding from the Canadian Institutes for Health Research (FDN-148453), a Brain Canada Platform Support Grant and a Canada First Research Excellence Fund award to BrainsCAN.

## Author information

### Contributions

R.M and R.S.M involved in conceptualization of paradigm. L.M.K developed the sequence for 9.4 T Bruker scanner. O.O developed the sequence for 3 T Siemens scanner. R.M and AE designed and performed the NHP experiments. R.M designed and collected the human data. R.M analyzed NHP and human data. R.S.M. wrote the simulation code. S.E performed the NHP surgeries. R.M and R.S.M wrote the paper with additional contributions from of L.M.K.

## Extended data figures

**Extended Data Fig. 1.**
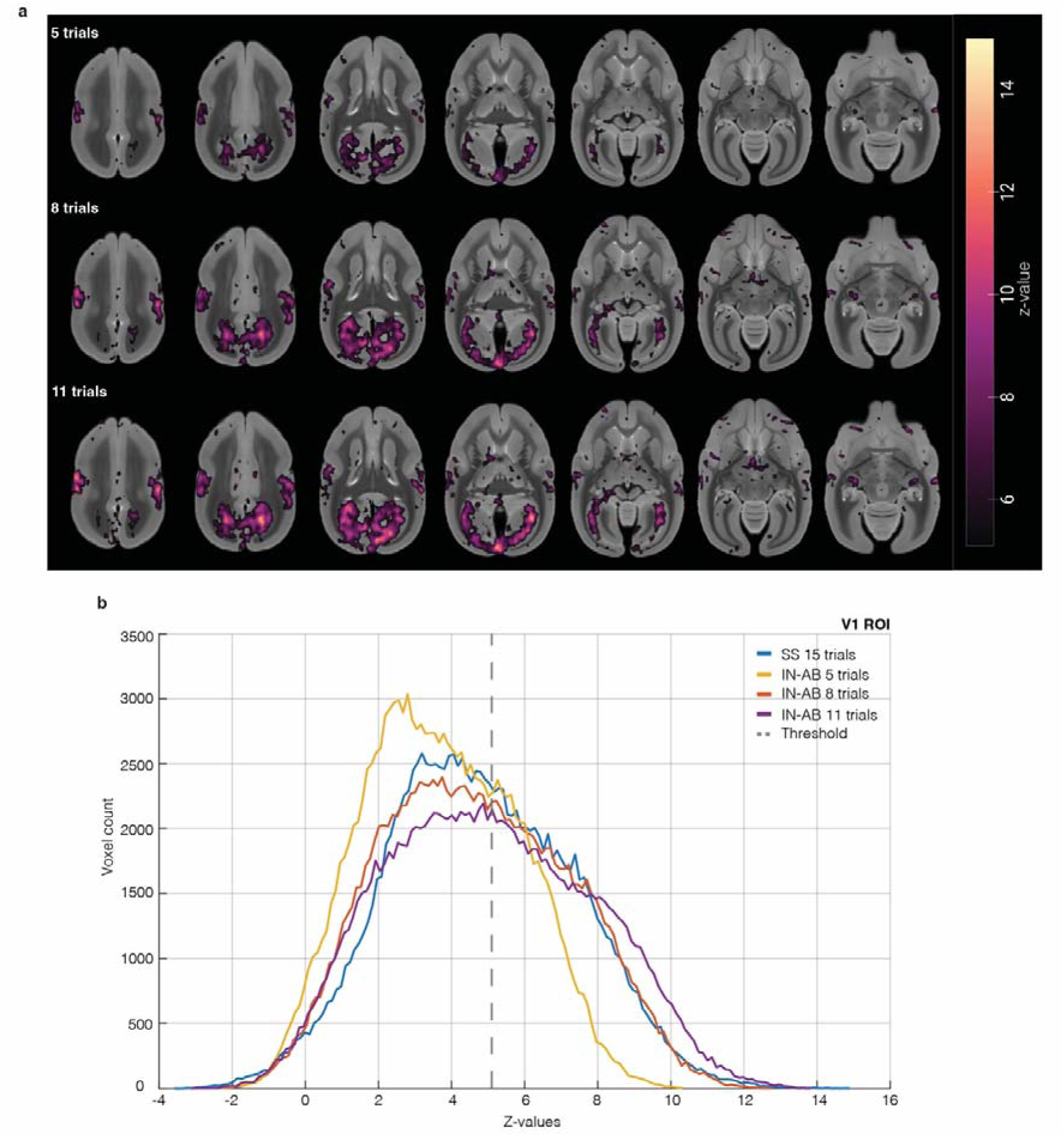
Reduced trails comparision (animal experiments at 9.4 T) **a.** Group activation map (n=3) for in AB visual stimulus experiment using SASS paradigm with different number of trials per run First row shows group activation from 5 stimulus trials per run (160 volumes), second row with 8 stimulus trials per run (256 volumes) and third row showing 11 trials per run (352 volumes - matching the acquisition time with SS paradigm). **b.** Average voxel count at each z-values for VI ROI with different reduced number of trials from SASS IN-AB paradigm with SS paradigm voxel count for 15 trials (480 volumes).

**Extended Data Fig. 2.**
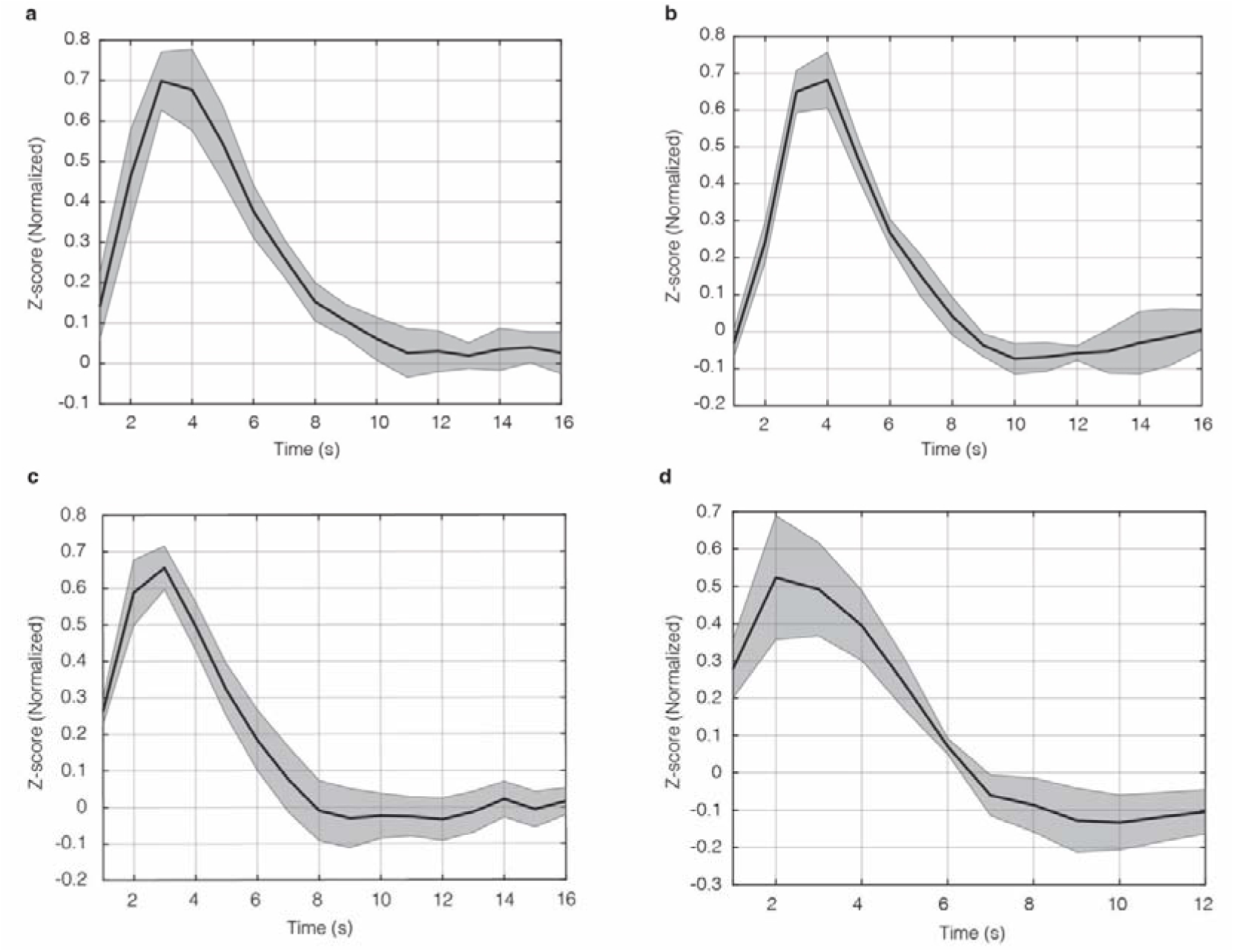
Mean of task active voxels across task AB/reglon (animal experiments at 9.4 T). **a,** SASS-paradigm In-AB visual stimuli BOLD response as mean z-score (normalized to maximum value) of task ABs across all subject runs (n= 3, P< 0.025, z-value> 5.1,). **b,** SS paradigm Visual stimuli BOLD response as mean z-score (normalized to maximum value) of task regions across all subject runs (n= 3, P< 0.025, z-value> 5.1). **c,** SASS paradigm pre-AB visual stimuli BOLD response as mean z-score (normalized to maximum value) of task ABs across all subject runs (n= 3, P< 0.025, z-value> 5.1). **d,** SASS paradigm pre-AB auditory stimuli BOLD response as mean z-score (normalized) of task ABs across all subjects runs (n=2, P< 0.025, z-value> 3.1).

**Supplementary Figure. 3.**
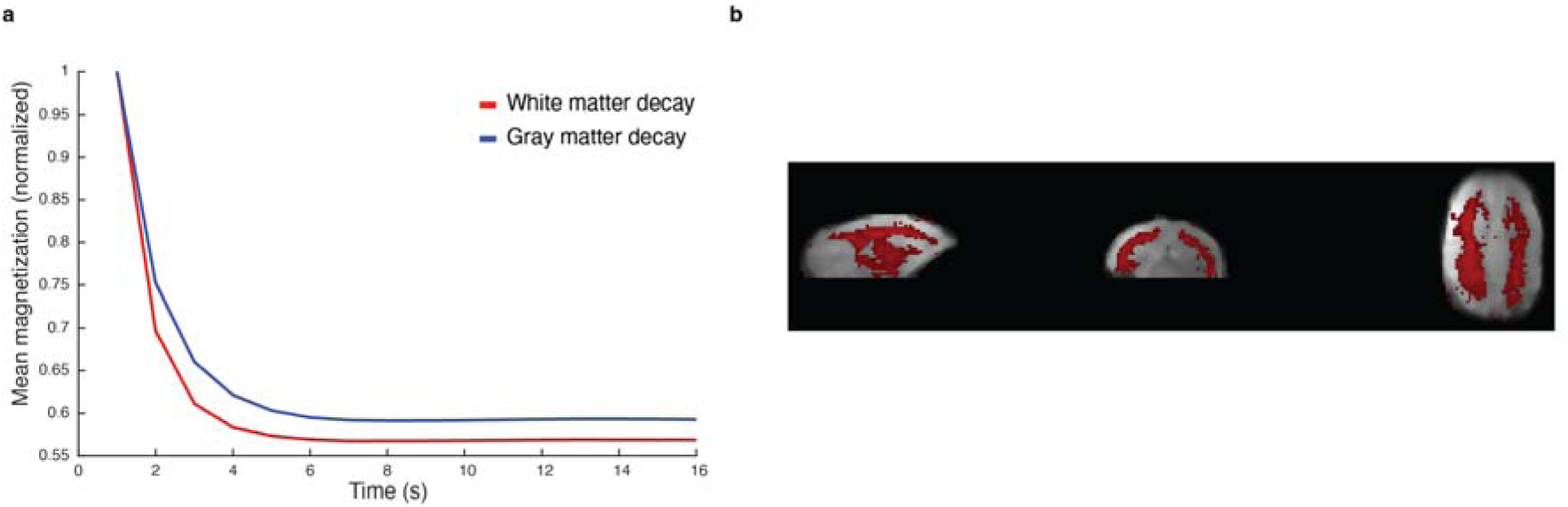
Segmentation using T1 decays. **a,** Mean magnetization decays of gray matter and white matter voxels across all ABs in a run. normalized to the maximum value. **b,** ICA component mask for white matter voxels from raw timeseries of one run.

**Extended data. Fig. 4.**
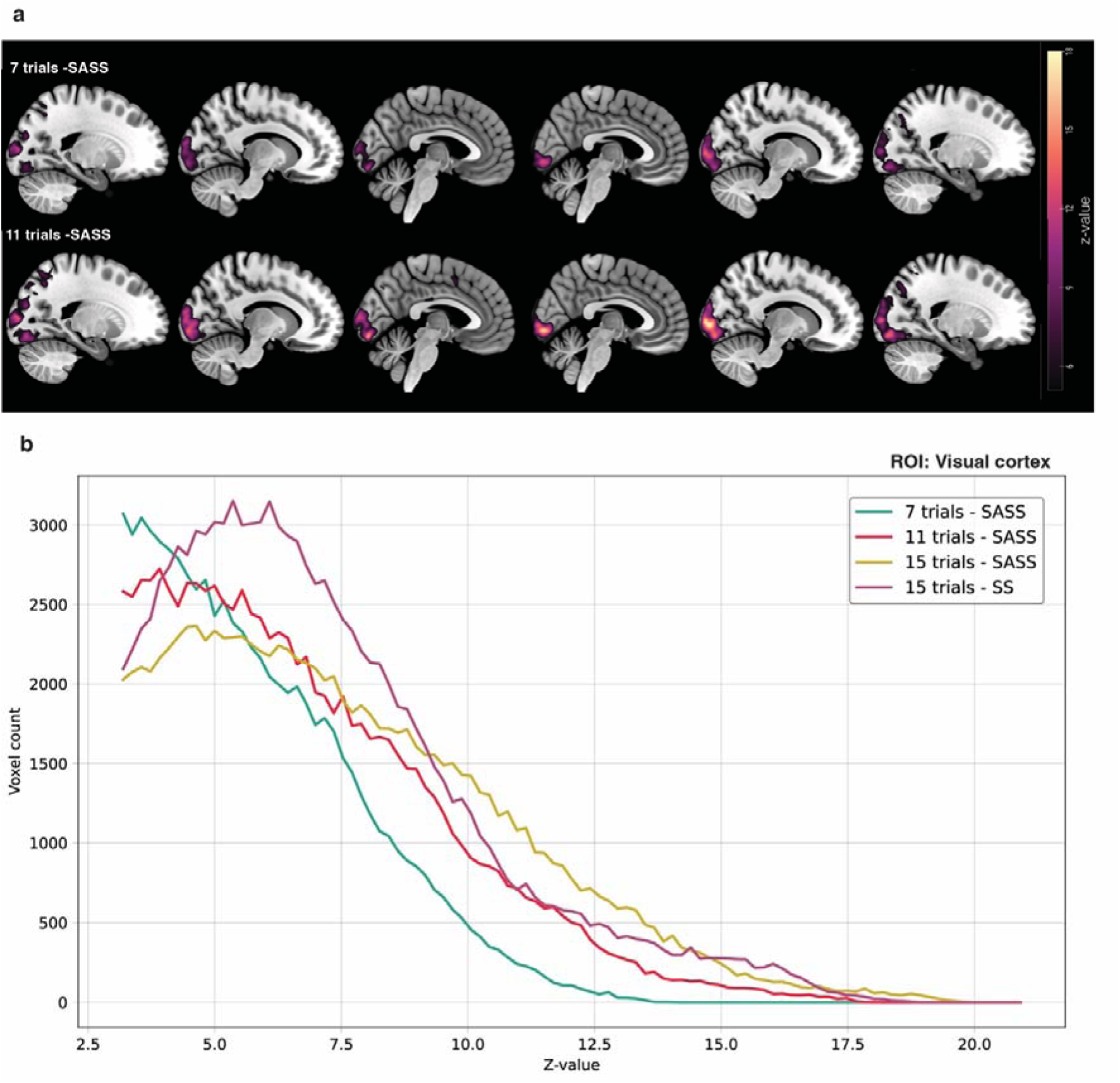
Reduced stimulus trials at human 3 T. **a,** SASS z-value maps from human experiments reduced number of stimulus trails First row showing z-value maps with 7 trials (240 volumes) and second row showing 11 trials (352 volumes) matching the scan time of SS experiment (15 trails). **b,** Voxel count at different z-values from visual cortex ROI for different SASS trial counts and SS experiment (n=3, P<0.05, z-value>5.1).

**Extended data. Fig. 5.**
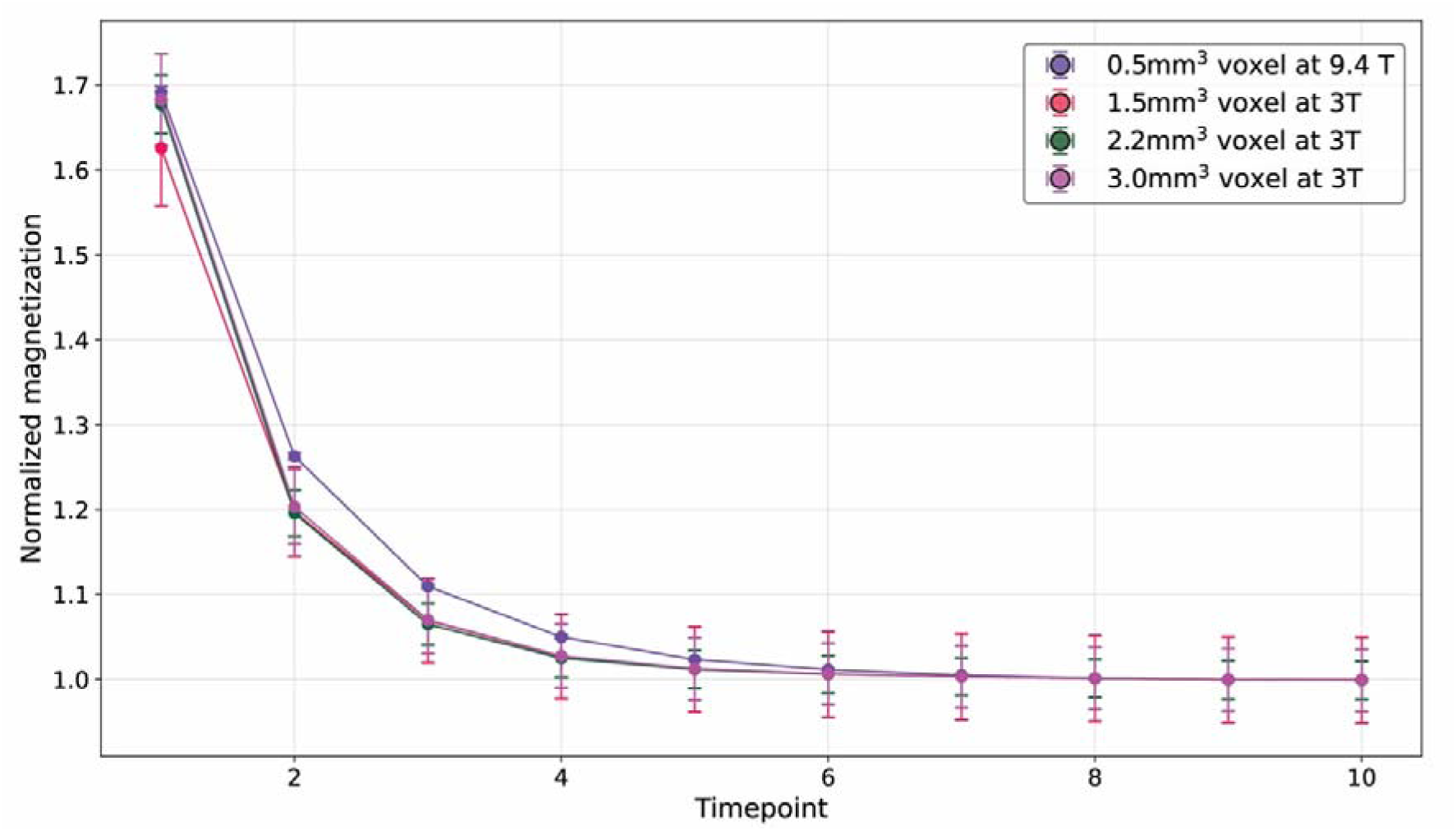
Grey matter magnetization decays. Magnetization across a 10s long AB at different voxel resolutions for 3 T human and comparative 9.4 T animal data. All voxel resolutions at 3 T were acquired with TR, TE and flip angle held constant (n=3) Comparative data from 9.4 T animal data shows a slower approach to steady state due to longer T1 in marmoset monkeys.

